# Unlocking the Potential of Low Quality Total RNA-seq Data: A Stepwise Mapping Approach for Improved Quantitative Analyses

**DOI:** 10.1101/2024.09.30.615750

**Authors:** Jiwoon Lee, JungSoo Gim

## Abstract

High-throughput sequencing assays face persistent challenges when analyzing low-quality RNAs, often assessed by the RNA integrity number (RIN). Current preprocessing methods and pipelines designed for mRNA-seq presume high-quality RNAs, overlooking the nuanced complexities arising from degraded transcripts in low-quality samples. This study questions the applicability of standard analysis pipelines, especially when sequencing low-quality total RNAs, which are sometimes the sole recourse for specific inquiries. To address this, we conducted a comprehensive investigation using large sequencing reads obtained from blood biospecimens with varying RINs. Introducing a novel mapping approach, termed ’stepwise mapping’, our systematic comparative analyses propose an optimal practice for total RNA-seq data analysis. Contrary to conventional mapping procedures, the ’stepwise mapping’ approach unveils additional transcriptome information, crucial for stable differential expression analysis, even with total RNA-seq data from specimens with relatively low RINs. Our method proves particularly valuable when analyzing limited specimens with low RNA quality.

## INTRODUCTION

Transcriptomic profiling has emerged as a valuable biomarker, providing indirect insights into an individual’s health status and the underlying biological mechanisms of diseases. Its active integration in the biomedical field is evident[1–4]. Particularly noteworthy are the surprising findings regarding the potential of transcriptomic biomarkers for early cancer diagnosis and risk prediction of degenerative diseases using blood samples[5–9]. These discoveries have significantly accelerated transcriptome research. With the growing importance of transcriptomes in disease studies, RNA-seq analysis of numerous clinical samples is underway. The non-invasiveness and ease of collecting blood samples, coupled with the significance of preventive medicine and early disease detection, have heightened the importance of transcriptome biomarkers. Nonetheless, challenges persist due to the inherent instability of transcripts, which leads to sample degradation and compromises sample quality in clinical research settings[10–13].

Transcriptomic research commences with the acquisition of high-quality RNA samples. Previous studies quantify sample quality by defining RIN (RNA integrity number), which reflects the degree of RNA degradation, and propose the threshold for sample quality analysis[14]. However, some issues in previous research remain unsolved. Firstly, the existing quality threshold primarily focuses on mRNAs [15–19], overlooking other transcriptome analyses. Secondly, limited analytic solutions are available for datasets where a specific RIN quality threshold cannot be met by the currently employed methods.

In the former cases, it is essential to verify the validity of applying the same quality standards used for mRNA-seq to the relatively more recent total RNA-seq. mRNA-seq frequently experiences 3’ mapping bias due to the experimental limitation when selecting mRNAs [15, 20, 21]. Moreover, the degradation of transcripts and subsequent quality deterioration lead to a larger mapping bias, which further affects subsequent analysis results such as differential expression analysis[15, 21–24]. While it is crucial to maintain sample quality in mRNA-seq, total RNA-seq employs a negative selection method that only removes rRNA, thus avoiding mapping bias caused by quality degradation[21, 22, 25, 26]. Therefore, it is necessary to reevaluate the applicability of the existing quality restrictions that have been conventionally applied.

The latter issue, generally leading to more critical problems, arisen from many experimental processes including the sample storage environment (temperature and pH), storage period, and post mortem interval (PMI) [16, 27–30]. In clinical, unlikely designed study, producing large-scale, high-quality data requires a considerable amount of resources and high costs[31, 32], and in some cases, only low quality samples are available due to the limited accessibility. As will be described in Methods, we observed such a particular example from our cohort study, where we conducted total RNA-seq analysis from approximately 300 blood biospecimens and 6 were filtered out for further analysis because they did not meet the commonly applied quality standard (RIN ≥ 6). Hence, there is a crucial need for an appropriate analysis method that can address the challenge effectively, in addition to meticulous sample collection and experimental processing.

In this study, we examine quality standards suitable for performing total RNA-seq for biomarker research and propose a stepwise mapping approach to handle inevitably obtained low-quality samples. To achieve the goal, we re-extracted 11 blood samples for total RNA-seq, including those that did not meet the quality criteria, resulting in 11 replicate pairs from the same individual with different RIN values. We then conducted a systematic comparison analysis to demonstrate the stability and differentiation of the proposed approach, offering a new opportunity for total RNA-seq analysis with samples that have been limitedly used due to quality constraints. Our study aims to address the issues of sample loss and associated costs, as well as the lack of data often encountered in clinical practice.

## RESULTS

### An Overall Workflow of The Proposed Approach Dealing with Low-Quality RNA Samples

An overall workflow is summarized in Figure 1. We utilised replicated sequencing data with different RIN for analysis. The RIN across the entire dataset ranged from 4 to 7.6, with differences in RIN between individual replicates varying from 0 to 3.2 (see supplementary Fig. S1). We employed a standard pipeline[33] for all preprocesses except for read mapping. To evaluate the results based on transcript quality and mapping performance, we conducted mapping using both the widely used standard method and a newly proposed method (see Method for further details). Our proposed method assumes that inherently short RNAs in total RNA-seq and degraded RNAs, which primarily contribute to low RIN in total RNA-seq, have been shortened while maintaining their original sequences. The key feature of our proposed method involves an additional step of realigning unmapped reads under adjusted conditions, following the same mapping procedure as the standard method. In contrast to the existing standard method, information of previously unmapped reads was obtained by performing additional mapping under looser conditions using a comprehensive reference genome containing assembly information of the entire genome region that is updated periodically. Subsequently, we quantified the expression levels of transcripts for each method, conducted several bioinformatic and statistical downstream analyses, and compared the results (Fig. 1). This comparison allowed us to demonstrate the effectiveness and utility of our proposed method, especially for samples previously excluded due to low RIN. The novel approach we propose has the potential to address issues related to sample loss, cost, and data scarcity, commonly encountered in actual clinical practice.

**Figure 1.**
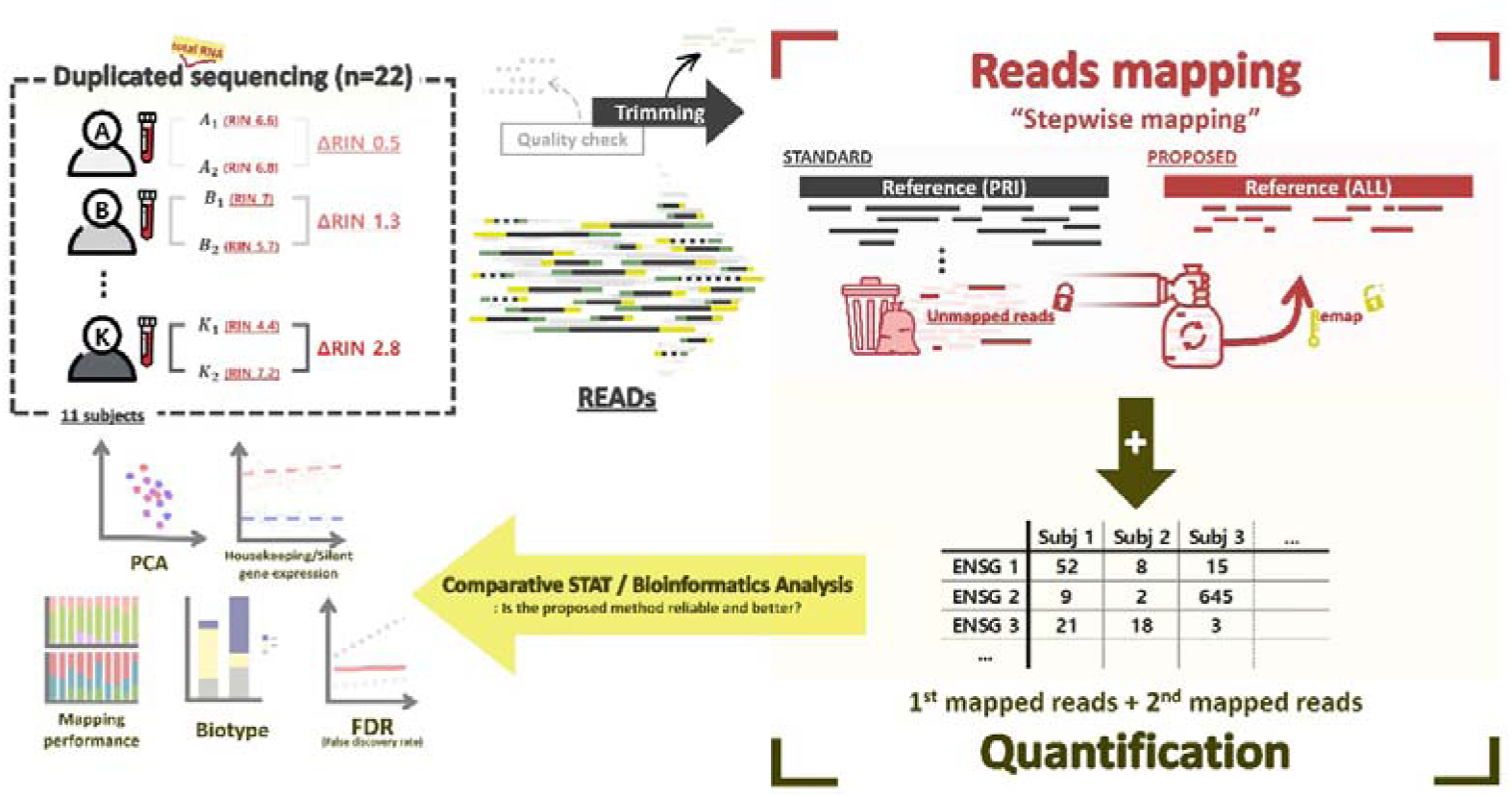
An Overall Workflow. Overall workflow for samples and studies. Blood buffy coat biopsy samples from 11 individuals were repeatedly sequenced to obtain technically replicated transcriptome data with different batches and RINs. Raw sequencing data were used for analysis after undergoing general transcriptome data preprocessing processes such as QC and trimming. Mapping was performed twice using a standard method that has been widely used and a new proposed method. The proposed method uses different reference genomes and mapping mechanisms to obtain more read information by remapping reads that were not mapped in the standard method under looser conditions. Afterwards, the estimated expression of genes was quantified and the stability and differentiation of the proposed method were validated through systematic comparative analysis.

### Enhanced Mapping Performance and Genomic Features Annotation

To begin, we conducted a comparison of mapping performance between the proposed method and the standard method, along with an analysis of genomic features associated with the mapped reads. Notably, the proposed method exhibited a substantial increase in remapping success for the previously unmapped reads, reducing the proportion of unmapped reads from 8.3% (SD: 7.02%) in the standard method to a mere 0.34% (SD: 0.53%). Furthermore, while the standard method often encountered difficulties in mapping shorter reads, the proposed method effectively mapped most of these shorter reads to specific regions in the genome. Specifically, the percentage of single-mapped reads increased from 65.2% (SD: 0.07%) in the standard method to 68.3% (SD: 0.05%) in the proposed method, resulting in richer information on uniquely mapped reads (Fig. 2A-B). In our more detailed comparison, we analysed and annotated the genomic features of the mapped reads (see Figure 2 C-D). The proposed method demonstrated significant improvement as 46.7% (SD: 15.13%) of reads that could not be mapped in the standard method were successfully remapped to exonic regions. Additionally, we observed further remapping to intronic (23.2%, SD: 4.7%) and intergenic (17.2%, SD: 7.4%) regions in the proposed method (Fig. 2E).

**Figure 2.**
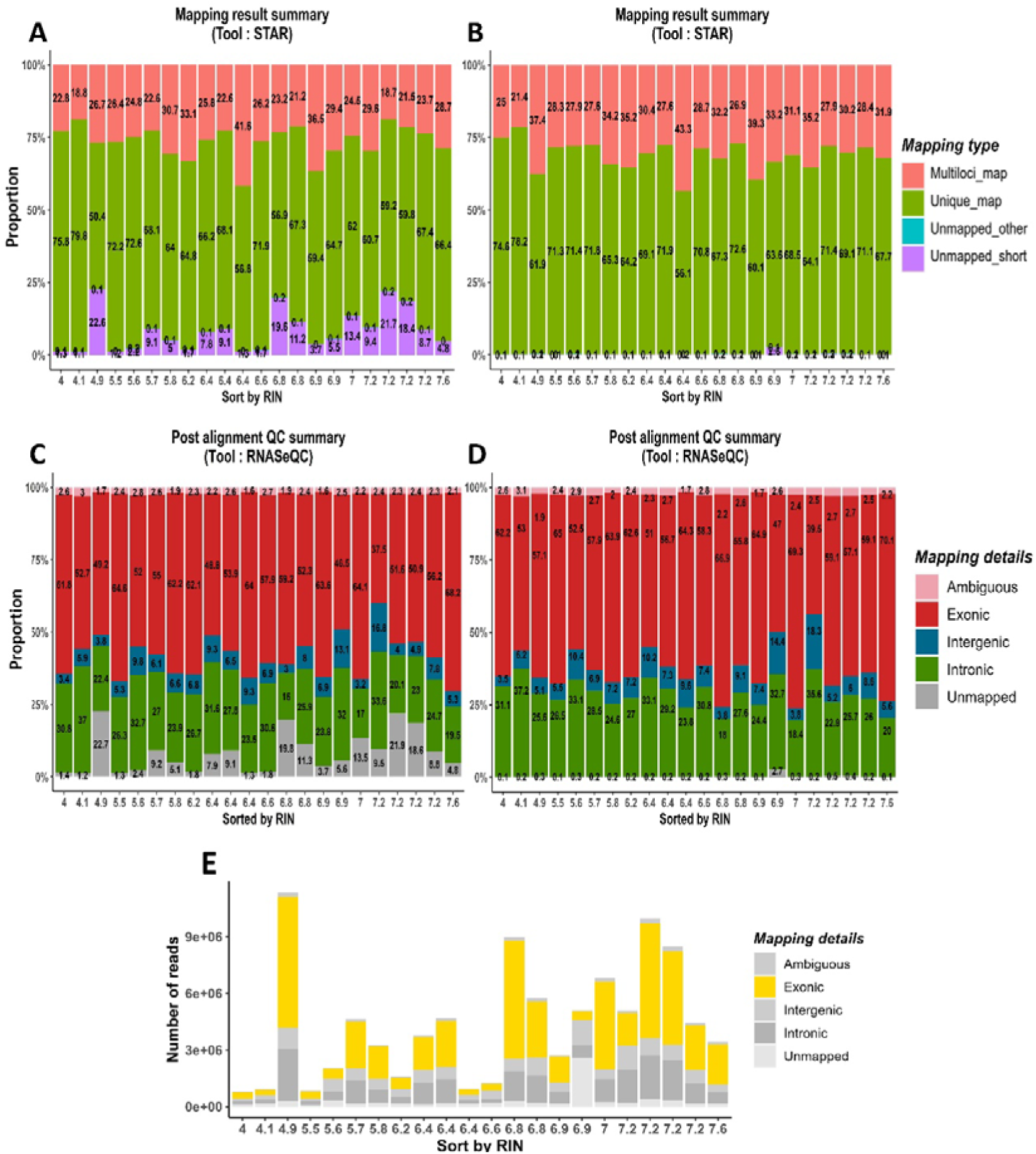
Mapping performance of standard method and proposed method. (A) Mapping performance of the standard method, (B) Mapping performance of the proposed method, (C) Genetic characteristics of reads mapped by the standard method, (D) Genetic characteristics of reads mapped by the proposed method, (E) Genetic characteristics of additional mapped reads only in the proposed method. The x-axis shows RINs sorted in ascending order, and the y-axis shows the number or ratio of reads. Mapping performance was provided by STAR, a read aligner, and genetic characteristics of mapped reads were provided by RNASeQC, a post alignment tool.

Regarding the correlation between sample quality and mapping performance, only the standard method displayed a significant positive correlation between the percentage of unmapped reads caused by the quality difference (beta regression, P < 0.05). There were no significant correlations between proportions of exonic reads, intergenic reads, ambiguous reads, and quality, but intronic reads showed a significant negative correlation (see supplementary Table S1).

Despite encountering low-quality samples with considerable rRNA degradation, rRNA depletion performed well in both methods. Consequently, the ratio of rRNA reads did not follow a distinct pattern across different qualities, with rRNA reads generally sparse across all qualities. While the proposed method decreased the rRNA ratio due to mapping more reads, the pattern remained similar to the standard method (Fig. 3). Then, comparing read mapping bias, we observed that the standard method exhibited more biased mapping, with an average of 46% (SD: 2.4%), while the proposed method achieved a more balanced mapping of reads at 51% (SD: 4%) (Fig. 3). This comparison demonstrated that the proposed method successfully remapped a greater number of reads to exonic regions, providing additional information that was not attainable using the standard method. As a result of this, the proposed method allows for more detailed transcriptome profiling in the downstream analysis.

**Figure 3.**
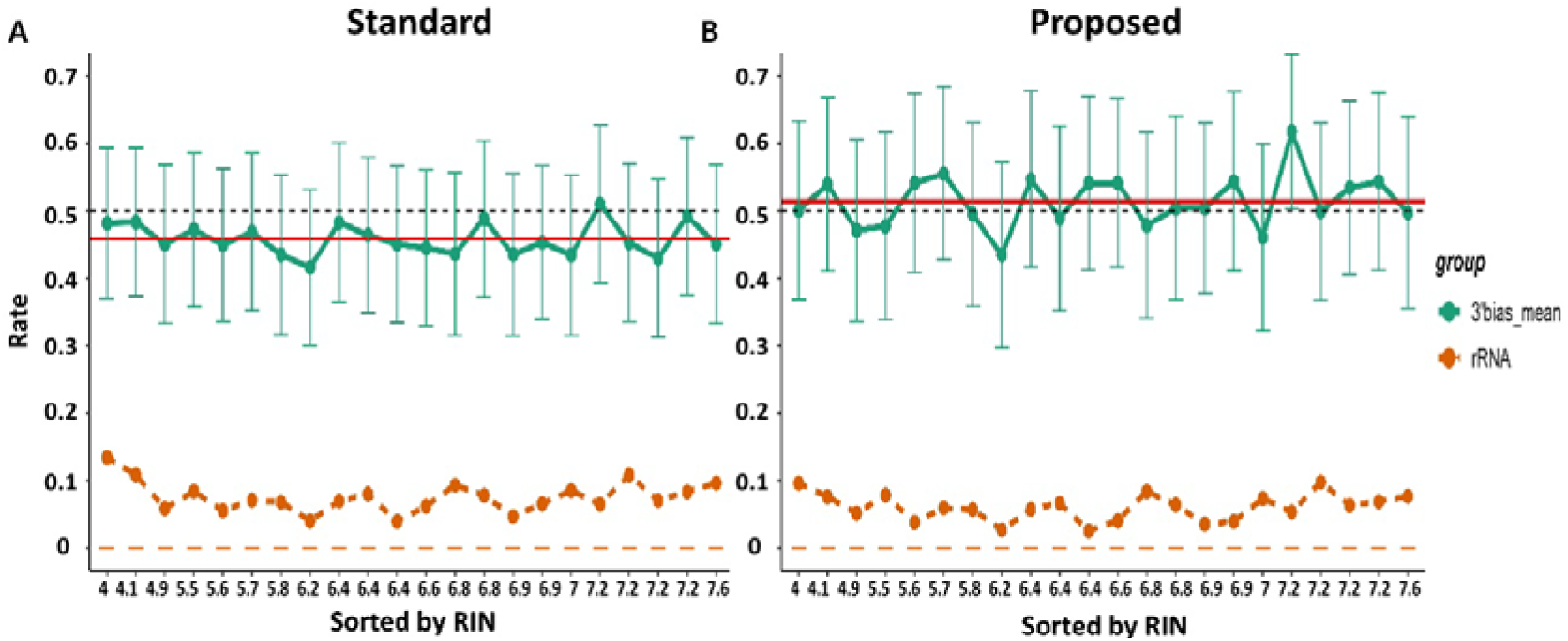
Mapping bias and rRNA contamination of standard method and proposed method. Comparison of 3’ mapping bias and rRNA contamination levels between standard and proposed methods. (A) Standard method, (B) Proposed method. The black dotted line located at 0.5 represents the average mapping coverage of genes that can be obtained when sequencing reads are ideally distributed uniformly across all genome regions. Exceeding this line means that the reads are mapped non-uniformly. Each green dot represents the average degree of bias for the sample, the error bar represents the 95% confidence interval, and the red line represents the average of the average mapping bias for the entire sample. The orange line at the bottom represents the rRNA contamination rate in the sample. Although the degree of rRNA contamination was reduced more in the proposed method (A) than in the standard method (B), the overall pattern is similar for the two methods because the proposed method detected more reads and the scale was reduced.

### Effect of Mapping Methods on Gene Expression Across Sample Quality Variation

To assess the impact of the mapping method and the sample quality on gene expression, we conducted analyses at the gene expression level. The proposed method detected the expression of 49,950 genes, while the standard method detected 48,986 genes, representing approximately 1,000 more genes detected in the proposed method. Additionally, we performed principal component analysis (PCA) to compare the overall expression patterns of all genes (Fig. 4A-B). Remarkably, the expression patterns of genes mapped using both the standard and proposed methods exhibited similarity without noticeable differences. Moreover, gene expression did not show any significant differences regardless of the sample quality.

**Figure 4.**
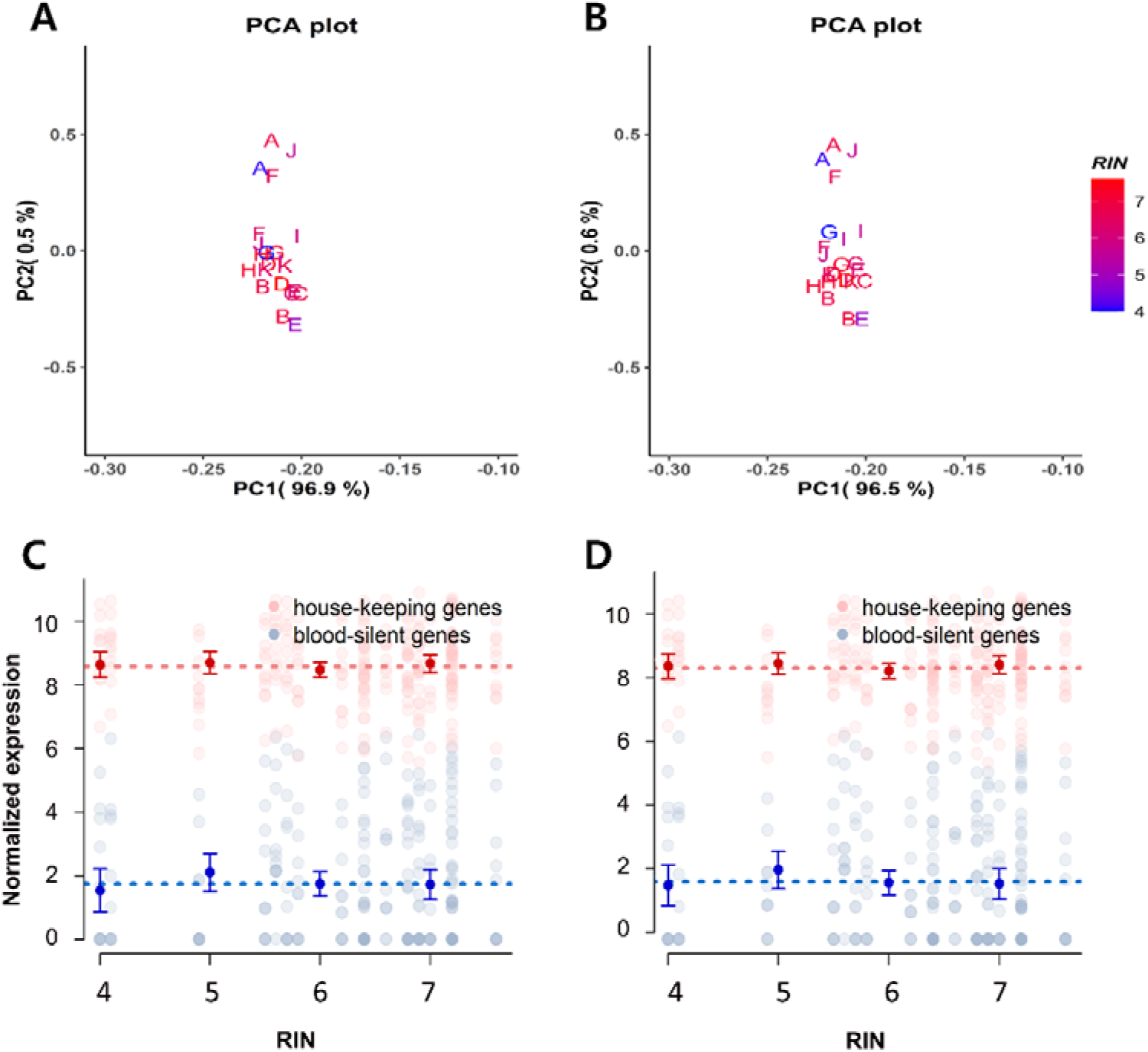
Expression profiling. (A) PCA plot of the standard method, (B) PCA plot of the proposed method. To reduce the dimensionality of overall gene expression for each sample, the two PC values with the highest explanatory power were used, and they explain more than 97% of the raw data. Each coordinate represents PC1 and PC2 values, and sample individuals were coded and identified by alphabet, and the same alphabet represents replicates. The RIN of the sample was coded in gradient color. (C) Expression of house-keeping/blood-silent genes using the standard method and (D) expression of the proposed method. The x-axis represents RIN and the y-axis represents normalized expression. Expression of 11 house-keeping genes and 11 blood-silent genes. Each dotted line represents the average expression of 11 genes for the entire sample by gene group, and the error bar represents the average and SD value of the average for each categorized RIN.

Most samples with varying quality (ΔRIN 0 to 3.2) demonstrated similar expressions, with their positions closely clustered or overlapping in the PC plot. These results indicated that not only the overall sample set but also the majority of replicates exhibited consistent gene expression regardless of the quality differences.

We also assessed the reproducibility of expression for both housekeeping genes and blood-silent genes across different RIN values. As can be seen in Fig. 4C-D, our data yielded average expression levels that were high for housekeeping genes and low for blood-silent genes, aligning with the known patterns from the GTEx project [52,53]. Additional analysis comparing the confidence intervals of expression by category, while categorizing quality, revealed that the confidence intervals remained constant and did not deviate significantly from each mean in both mapping methods. Furthermore, regardless of variation in sample quality as measured by the RIN scale, the expression levels of both housekeeping genes and blood-silent genes remained consistent. This demonstrates that changes in sample quality did not correlate with variation in gene expression in both methods (Fig. 4C-D). Based on these findings, which suggest that the proposed method can be reliably applied to samples with diverse quality levels without affecting the expression analysis outcomes, we conducted further analysis to assess the validity of the proposed method compared to the existing method.

### Comparative Analysis of Differential Expression Using Duplicated Samples

The sequential comparative analysis between the proposed method and the standard method aimed to showcase the differences and the advantages of the proposed method. Differential expression (DE) analysis was performed on a study sample of 22 replicates (duplicates from 11 subjects), and differentially expressed genes (DEGs) were detected from each pair of replicates. DEGs detected in the analysis are interpreted as errors resulting from either the limitation of preprocessing method or the variability of sample quality, providing a critical measure of method performance.

Figure 5 clearly demonstrates that the proposed method outperforms the standard method in controlling false discoveries, particularly evident when dealing with larger differences in sample quality, including low-quality samples. In the standard method, as the quality difference between samples increased, the rate of false discoveries steeply rose, exceeding statistical standards across significance levels (Fig. 5A). Conversely, the proposed method maintained better control over false discoveries, with a significantly lower percentage of samples exceeding false discovery rate (FDR) at all significance levels (Fig. 5B). Notably, in the proposed method, the regression line describing the relationship between the RIN difference between samples and the FDR did not surpass the significance level of 5% and showed a gradual increase as the RIN difference increased (red solid line in Fig. 5B). Thus, in the proposed method, FDR remained relatively stable despite the increase in the quality difference between samples. Conversely, in the standard method, the regression line exceeded the significance level of 5% even when there was no difference in qualities between samples, and as the RIN difference between samples reached 3, it approached 10% (red solid line in Fig. 5A). Furthermore, when using the relatively less conservative significance level of 0.1, the proposed method effectively controlled FDR in all quality differences, as the regression line did not exceed the threshold of 10% (dark grey dotted line in Fig. 5B). However, in the standard method, the regression line increased steeply and exceeded, indicating that FDR was not controlled.

**Figure 5.**
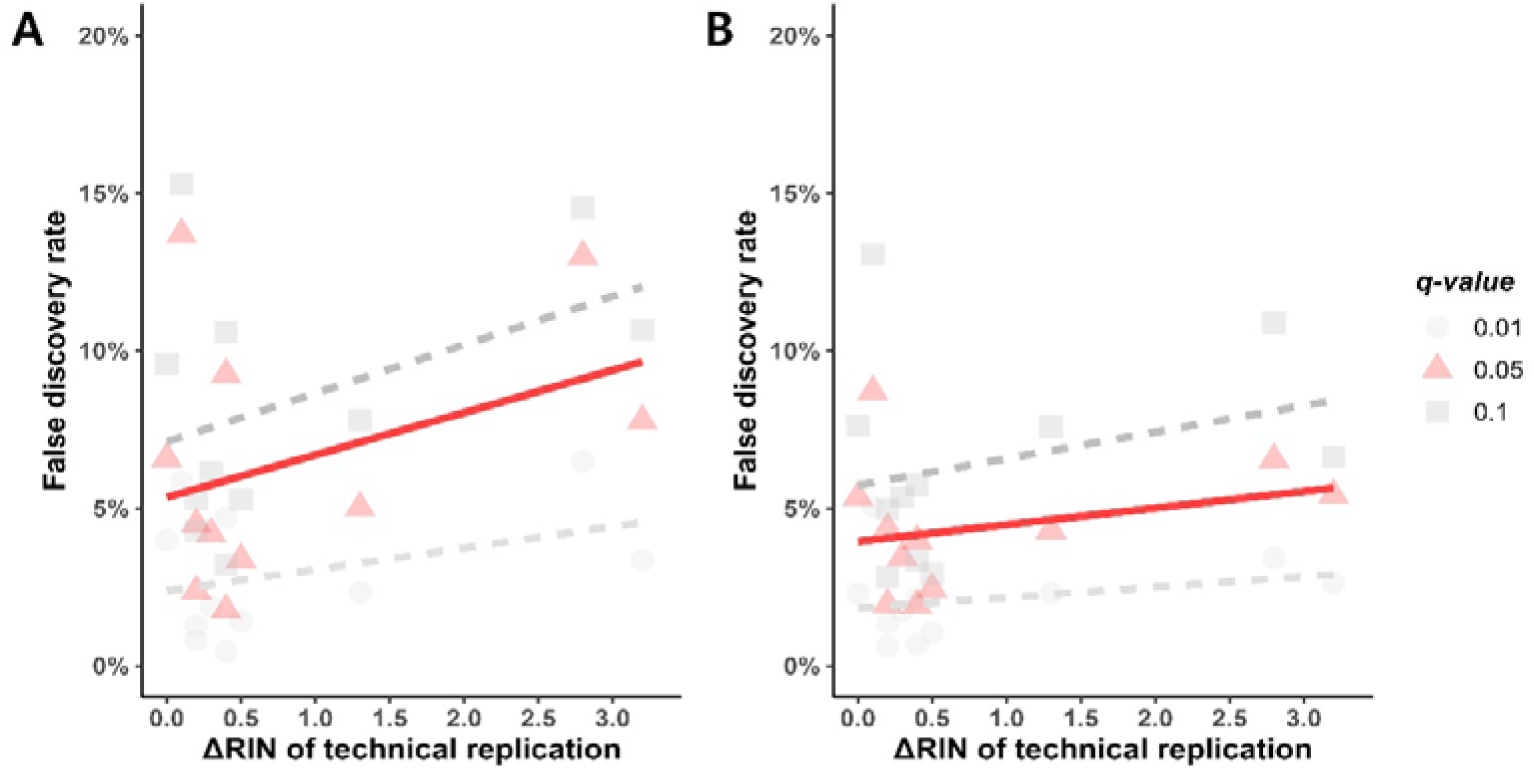
FDR control. The proportion of falsely discovered DEGs and their degree of statistical control (A) in the standard approach and (B) in the proposed approach. X-axis represents the difference in RIN between replicates, and the y-axis represents the FDR as the proportion of DEGs detected in the analysis after multiple comparison correction (Benjamini-Hochberg procedure). DEGs between replicates were defined based on the most commonly used statistical significance levels of 0.01, 0.05, and 0.1, respectively, and the results by significance level were distinguished by different colors and shapes.

Based on these results, we confirmed that gene expression quantification using the proposed method effectively controlled the inherent variability in transcripts statistically and further demonstrated the performance, even in data including low-quality samples.

### Biotype of newly mapped reads

We conducted additional comparative analyses to discern the individual effects of using different reference genomes and mapping strategies in the proposed approach. To ensure a fair comparison, we employed a standard reference genome instead of the one utilized in the proposed method. However, we established a ‘positive control’ comparison group by applying two step mapping strategy. Subsequently, we compared the biotypes of the additional identified genes using the proposed method, standard method, and the positive control.

To investigate the effects of additional mapping, we compared the standard approach with the positive control, which used the same standard reference genome but a different mapping strategy. Not surprisingly, all genes identified by the standard method were also detected in the positive control, but the positive control group identified approximately 1,116 genes (Fig. S2). Conversely, to examine the effects of reference genomes, we compared the proposed method with the positive control group, which employed different reference genomes but the same proposed mapping strategy. Interestingly, we observed that each reference genome led to the specific detection of expressed genes. The positive control group, using the standard reference genome, detected 1,645 genes, whereas the proposed method detected 1,483 genes, with more than 53.1% of the genes specifically detected by the proposed method originating from protein-coding regions. The majority of the genes specifically detected only in the positive control group were pseudogenes (31.7%), lncRNAs (24.7%), and protein-coding genes (23%) (Fig. 6 and Tables S2-3).

**Figure 6.**
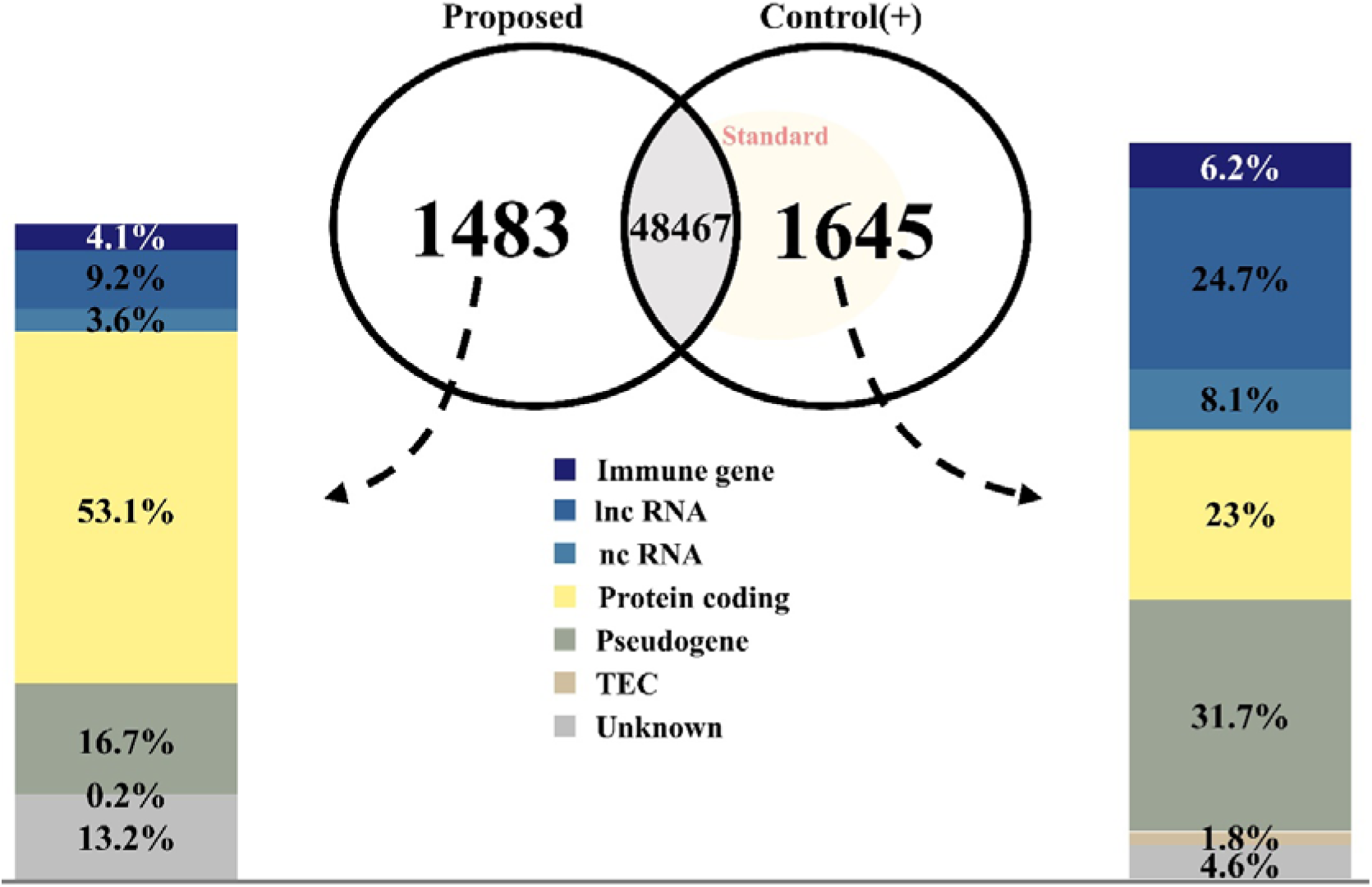
Biotype Comparison. As a result of comparing biotypes between the proposed method, the standard method, and the positive control mapping method, the positive control used the standard reference genome and the proposed mapping method (mechanism). All genes detectable in the standard method can be detected in positive control mapping, there are 1,645 genes whose expression is specifically detected only in positive control mapping, and 1,483 genes whose expression is specifically detected only in the proposed method. The biotypes of genes specifically detected for each method are shown in a bar plot, and more than half of the genes detected only in the proposed method are protein-coding genes.

Comparing the number of genes quantified and merged by all approaches, we found that the proposed approach, the standard approach, and the positive control respectively quantified 49,950 genes, 48,986 genes, and 50,112 genes (Fig. S2). Notably, both the proposed and standard approaches identified a considerable number of genes, with 1,588 genes specifically quantified by the standard method and 2,550 genes specifically quantified by the proposed method (Fig. S2). Among the genes specifically quantified by the proposed method, more than 33% (853 genes) were protein-coding genes, with additional representation from lncRNAs (10.4%), ncRNAs (10.5%), and pseudogenes (33.6%); in contrast, the genes specifically detected by the standard method comprised 23.5% protein-coding genes (373 genes), along with lncRNAs (25.3%), ncRNAs (7.9%), and pseudogenes (30.7%) (Fig. S3 and Tables S4-5). Despite total RNA-seq targeting all RNA types, the proposed approach detected a larger number of genes, particularly from protein-coding regions. Despite total RNA-seq targeting all RNA types, the proposed approach detected a higher proportion of genes from protein-coding regions compared to the standard approach. Taken altogether, both the different reference genomes and mapping methods used in the proposed approach contribute to enhancing the performance of transcriptome analysis.

## DISCUSSION

The degradation of transcripts within living cells bears significant implications, as it regulates gene expression, thereby impacting all biological responses[34–36]. Moreover, as it directly influences sample quality, a pre-sequencing quality check of samples becomes imperative[18, 33, 37]. Traditionally, the quality of transcripts is ascertained based on the size and concentration of two types of rRNA (18S, 28S) in the sample, alongside various other values and their relative ratios[14]. In mRNA-seq, one of the most common transcriptome analyses, poly A selection is employed to selectively extract mRNA, resulting in 3’ mapping bias due to degradation and consequent quality deterioration. Given that mapping bias directly affects gene expression quantification and subsequent analyses, sample quality limitations are inevitable in mRNA-seq. However, total RNA-seq employs negative selection, removing only rRNA through rRNA depletion, thereby avoiding mapping bias caused by quality degradation[15, 20, 22, 25]. Furthermore, in most RNA-seq experiments, high-quality samples are randomly fragmented, and subsequent size selection processes being employed to use only fragments of appropriate sizes for sequencing[38–40]. Thus, applying quality limitation based on transcript degradation lengths to total RNA-seq, which includes RNAs of various lengths, would only lead to resource wastage.

For these reasons, this study utilized samples with diverse qualities without bias. We proposed a mapping approach suitable for total RNA-seq analysis, incorporating lower quality samples than those used in previous studies’ quality standard. The validity and stability of the proposed approach were validated through systematic comparative analyses. The mapping with the proposed approach revealed that the majority of reads that could not be mapped in the standard approach were evenly mapped to exonic, intronic, and intergenic regions in the genome, providing more balanced expression information. The expression profiling and expression reproducibility verification between the standard and proposed approaches demonstrated consistent patterns, confirming the stability of the proposed approach. Additionally, the proposed approach could quantify the expression of about 1,483 more genes than the standard approach, and about a half (53.1%) of these additionally quantified genes were protein-coding genes. DE analysis of the quantified genes, considering RIN differences between samples, showed that the proposed approach effectively controlled false discoveries, even with increasing RIN differences.

Furthermore, we investigated whether sample quality influenced the analysis results. The results showed that sample quality did not significantly affect read mapping performance or expression quantification in both approaches. Particularly, analysis of the overall expression pattern among samples using PCA demonstrated that gene expression remained consistent even with a RIN difference of 3 or more, showing similar pattern. Additionally, the reproducibility of previously studied house-keeping genes and blood-silent genes, showed consistent expression levels regardless of RIN, confirming that sample quality (if RIN is larger than 4) did not impact downstream data preprocessing and analysis.

Our results indicated that the proposed approach outperformed the standard approach, showing improved performance and statistical outcomes. Nonetheless, caution is warranted when applying the proposed approach to samples with extensive genetic mutations stemming from structural damage to DNA, such as those found in cancer samples[41, 42]. The primary process of the proposed approach, the inclusion of secondary mapping, involves adjustments to match read sequences to the reference genome more loosely, thereby enabling the mapping of RNA of various lengths in total RNA samples with small RIN. While this adjustment facilitates mapping in samples with diverse RNA lengths, it may lead to inaccuracies in samples with unspecified genetic variations, potentially causing confusion in the results.

## METHODS & MATERIALS

### Study Sample and Consent

A cohort study, a separate research project from this work, conducted total RNA-seq on blood samples from approximately 400 subjects. All subjects enrolled in the study provides written and informed consent in accordance with the Institutional Review Board of Chosun University Hospital (IRB No. 2-1041055-AB-N-01-2020-37), Gwangju, Republic of Korea. We found samples did not meet the quality standard of for RNA sequencing (RIN 6 or higher). To obtain high-quality sample, we re-extracted intact samples and performed total RNA-seq on six blood samples filtered from a larger cohort study that did not meet quality criteria. Additionally, six samples were randomly re-selected from subjects who met the criteria, resulting in a total of 12 replicates pairs. One subject was excluded due to difficulty in obtaining consent for blood re-sampling, leaving us with a final set of 11 replicates pairs (n=22) for total RNA-seq data generation. This resulted in 11 replicate pairs with different RIN from the same person. Sequencing was conducted using the rRNA-depletion method, which targets all transcripts excluding ribosomal RNA (rRNA). The RIN value of the study sample (n=22) ranged from 4 to 7.6 (mean: approximately 6.3, median: 6.5).

### Total RNA sequencing

Total RNA Sequencing using Illumina Truseq Stranded Total RNA Library Kit. RNA was extracted from buffy coat using RiboPure™ RNA Purification Kit (Ambion) according to the manufacturer’s recommended procedures. Following total RNA purification, the sample was treated with DNase using DNA-Free kit (Ambion) to eliminate potential DNA contamination that could have interfered with the result interpretation. RNA purity was determined by assaying 1 µl of total RNA extract on a NanoDrop8000 spectrophotometer. Total RNA integrity was assessed using an Agilent Technologies 2100 Bioanalyzer, with an RNA Integrity Number (RIN) value assigned. Total RNA sequencing libraries were prepared according to the manufacturer’s instructions (Truseq Stranded Total RNA Library Human/Mouse/Rat kit, Cat # 20020597, Illumina). A total of 400 ng of total RNA was subjected to ribosomal RNA depletion using biotinylated probes that selectively bind rRNA species. Following purification, the rRNA-depleted total RNA was fragmented into small pieces using divalent cations under elevated temperature. The cleaved RNA fragments were copied into first-strand cDNA using reverse transcriptase and random primers, followed by second-strand cDNA synthesis using DNA Polymerase I and RNase H. These cDNA fragments then had a single ’A’ base added and underwent subsequent ligation of the adapter. The products were purified and enriched with PCR to create the final cDNA library. The quality of the amplified libraries was verified by automated electrophoresis (Tapestation, Agilent). After qPCR using Kapa illumina library quantification Kit, libraries that were index-tagged were combined in equimolar amounts in the pool. RNA sequencing was performed using an Illumina NovaSeq 6000 system following provided protocols for 2×100 sequencing.

### Total RNA-seq Data Preprocessing

The transcriptome data were analyzed using the standard pipelines and tools for preprocessing sequencing data, which included sequencing quality assessment, adapter removal, annotation of mapped reads, and gene expression quantification. Sequencing quality assessment was conducted using *FastQC*[43], which involved evaluating sequencing base quality and adapter contamination to determine the need for adapter removal. Adapter removal was performed using *bbduk*, toolset of *BBtools* (`ref=adapter.fa, ktrim=r, k=23, mink=11, hdist=1, tpe, tbo`). Subsequent quality assessment was conducted to ensure more accurate results. Read mapping and gene expression quantification were performed using the *STAR* aligner[44], with the reference genome build being Human GRCh38 (gencode release version 38, https://ftp.ebi.ac.uk/pub/databases/gencode/Gencode_human/release_38/). For mapping, two types of reference genomes and two different mapping approaches were combined to compare the results between the standard and proposed approach. The standard approach used a typical reference genome containing primary assembly information for commonly used genes in transcriptome analysis (denoted as ‘PRI’) and followed the default options recommended by the aligner. In contrast, the proposed mapping approach utilised a comprehensive reference genome containing updating assembly information for all genes (denoted as ‘ALL’) and involved a two-step process with looser mapping options for reads that failed to map using the default options of the first standard mapping approach. The second mapping step in the proposed approach included additional options related to data input and output `(“--readFilesCommand zcat”, “--outReadsUnmapped Fastx”, “--outSAMtype BAM SortedByCoordinate”, “--quantMode GeneCounts”)`, and modified options to allow for looser mapping `(“--quantMode GeneCounts”, “--outFilterMatchNminOverLread 0.4”, “-- outFilterScoreMinOverLread 0.4”, “--outFilterMultimapNmax 20”)`. *RNA-SeQC*[45] was used to annotate genomic features of reads mapped by each method, obtaining the number of reads mapped to exonic, intronic, intergenic, and other regions using the same reference genome as used for mapping. Gene expression quantification was achieved by using the `“--quantMode GeneCounts`” option within the *STAR* aligner, generating expression counts equivalent to those obtained using *HTSeq-count*[46], a quantification tool. Depending on the downstream analysis objectives, genes with zero total sum of count across the entire sample were either excluded from analysis or all detected genes were retained for further analysis.

### Statistical and Bioinformatics Analysis

All statistical and comparative analyses, along with their results, were performed and generated using R/Rstudio[47, 48]. A downstream statistical analysis was conducted using *LPEseq*[49], a R package for DE analysis of non-replicated RNA-seq data. Gene expression was standardized using the normalization function “`LPE.normalise()”,` which employed log transformation considering library size. Most of the results were visualized using the *ggplot2*[50].

1. Principal Component Analysis (PCA): Principal component analysis (PCA) was performed to compare the overall expression structure of study sample. PCA was used to reduce the high-dimensional expression matrix of approximately 50,000 genes to low dimensions. The analysis utilised the R built-in function “`prcomp()`” to reduce gene expression per sample and used the top 2 PCs, explaining over 97% of the total gene expression, for comparison[51].
2. Differential Expression (DE) Analysis: For DE analysis between non-replicated duplicates from the same subject, *LPEseq* was used. To address multiple comparison issues, resulting p-values were adjusted using the Benjamini-Hochberg method for false discovery rate (FDR) control, and DEGs were selected based on adjusted p-values. Commonly used significance levels of 0.01, 0.05, and 0.1 were employed, and the proportions of DEGs selected at each significance level were used in the analysis to assess for FDR control.
3. Housekeeping and Blood-Silent Genes Analysis: To evaluate the stability of the proposed approach, the expression reproducibility of genes with known expression patterns was investigated. Genes with validated expression stability from previous studies were used, categorized into housekeeping genes and blood-silent genes based on their reported characteristics. Housekeeping genes are composed of genes that consistently maintain a certain level of expression across tissues as well as is blood[52], while blood-silent genes were selected from the GTEx project’s tissue-specific gene expression information for genes with low expression levels in blood (https://www.gtexportal.org/home/tissue/)[53]. Please refer to the supplementary data for the list of genes included in the analysis.
4. Biotype Annotation: To compare the biotypes of genes specifically quantified by each approach (proposed, standard, positive control), annotation was performed using the R package “EnsDb.Hsapiens.v86”. Detailed biotype classifications were categorized according to Ensembl’s biotype classification criteria for analysis (http://asia.ensembl.org/info/genome/genebuild/biotypes.html)[54].

## KEY POINTS

- The study questions standard preprocess pipelines’ suitability for sequencing low-quality total RNAs.
- Introduces ‘stepwise mapping’ for total RNA-seq data analysis.
- ‘Stepwise mapping’ uncovers additional transcriptome information crucial for stable differential expression analysis.
- Particularly valuable for analysing limited specimens with low RNA quality

## ACKNOWLEDGEMENTS

The authors thank the members and enrolled participants of the Gwangju Alzheimer’s & Related Dementias (GARD) study.

## DATA AVAILABILITY

The processed dataset and reproducible analysis code written in R supporting this article are available upon request or in the online links through https://github.com/jiwooo-on/total_RNAseq_pipeline/blob/main/Supplementary_files.zip or http://gimlab.online/data/total_RNAseq/Supplementary_files.zip.

## AUTHOR CONTRIBUTION

Jiwoon Lee: Scientific idea, programming and technical design, analysis and interpretation of results, manuscript preparation and revision.

Jungsoo Gim: Design of the study and experiment, scientific idea and technical design, interpretation of results, manuscript preparation and revision.

## BIOSKETCH

M.Sc. Jiwoon Lee has just started her professional career as a research associate at Chosun University since her two-year full-time Master’s program, which included coursework and research. During her tenure, she actively participated in a large cohort study, gaining extensive experience in RNA-seq data quality control and analysis.

Dr. JungSoo Gim holds a Ph.D. in bioinformatics from Seoul National University, complemented by an M.S. in theoretical particle physics and B.S. in chemistry. With a diverse interdisciplinary background, he received training in developing statistical methods and their application to complex diseases. He had held positions as a post-doctoral fellow and a research assistant professor at Seoul National University, as well as serving as a visiting scientist at Harvard University. Currently, Dr. Gim is a faculty member at Chosun University and co-directs the Gwangju Alzheimer’s & Related Dementias (GARD) Study. This study, supported by a number of grants including funding from the Korean government spanning over a decade and a $12,000,000 award from the US NIH/NIA, aims to address the pressing challenges of Alzheimer’s disease research.

