## supplementary for "Unlocking the Potential of Low Quality Total RNA-seq Data: A Stepwise Mapping Approach for Improved Quantitative Analyses"

**Supplementary Data (Tables and Figures)**

**Supplementary Table S1.** Result of correlation analysis between mapping performance and RIN

| beta-regression : Mapping performance ~ RIN (RNA quality) | | | | |
| --- | --- | --- | --- | --- |
| **Genome Features (%)** | **standard_β** | **standard_P** | **proposed_β** | **proposed_P** |
| Unmapped | 0.370 | 0.034 | 0.141 | 0.369 |
| Exonic | -0.015 | 0.822 | 0.016 | 0.807 |
| Intronic | -0.132 | 0.013 | -0.116 | 0.019 |
| Intergenic | 0.095 | 0.336 | 0.112 | 0.232 |
| Ambiguous | -0.044 | 0.212 | -0.031 | 0.358 |

**Table S2.** A detailed biotype list for comparison between mapping methods (Proposed – Control (+))

| PROPOSED-CONTROL(+) : Detected only in proposed method | | | |
| --- | --- | --- | --- |
| **Biotype** | **no.** | **Biotype** | **no.** |
| antisense | 26 | sense_intronic | 2 |
| IG_C_gene | 5 | snoRNA | 5 |
| IG_V_gene | 14 | snRNA | 6 |
| IG_V_pseudogene | 5 | TEC | 3 |
| lincRNA | 45 | TR_C_gene | 2 |
| miRNA | 1 | TR_J_gene | 5 |
| misc_RNA | 14 | TR_V_gene | 35 |
| polymorphic_pseudogene | 5 | TR_V_pseudogene | 5 |
| processed_pseudogene | 84 | transcribed_processed_pseudogene | 3 |
| processed_transcript | 62 | transcribed_unitary_pseudogene | 1 |
| protein_coding | 787 | transcribed_unprocessed_pseudogene | 48 |
| rRNA | 27 | unitary_pseudogene | 1 |
| scaRNA | 1 | unprocessed_pseudogene | 95 |

**Table S3.** A detailed biotype list for comparison between mapping methods (Proposed - Control (+))

| PROPOSED-CONTROL(+) : Detected only in CONTROL(+) | | | |
| --- | --- | --- | --- |
| **Biotype** | **no.** | **Biotype** | **no.** |
| 3prime_overlapping_ncRNA | 1 | protein_coding | 378 |
| antisense | 168 | rRNA | 8 |
| IG_C_gene | 2 | sense_intronic | 24 |
| IG_C_pseudogene | 2 | sense_overlapping | 4 |
| IG_D_gene | 21 | snoRNA | 12 |
| IG_J_gene | 6 | snRNA | 31 |
| IG_J_pseudogene | 3 | TEC | 29 |
| IG_pseudogene | 1 | TR_J_gene | 3 |
| IG_V_gene | 53 | TR_V_gene | 17 |
| IG_V_pseudogene | 60 | TR_V_pseudogene | 6 |
| lincRNA | 195 | transcribed_processed_pseudogene | 10 |
| miRNA | 45 | transcribed_unitary_pseudogene | 3 |
| misc_RNA | 38 | transcribed_unprocessed_pseudogene | 34 |
| polymorphic_pseudogene | 10 | unitary_pseudogene | 4 |
| processed_pseudogene | 224 | unprocessed_pseudogene | 164 |
| processed_transcript | 14 |  |  |

**Table S4.** A detailed biotype list for comparison between mapping methods (Standard – Proposed)

| STANDARD-PROPOSED: Detected only in the standard method | | | |
| --- | --- | --- | --- |
| **Biotype** | **no.** | **Biotype** | **no.** |
| 3prime_overlapping_ncRNA | 1 | protein_coding | 373 |
| antisense | 168 | rRNA | 7 |
| IG_C_gene | 2 | sense_intronic | 24 |
| IG_C_pseudogene | 2 | sense_overlapping | 4 |
| IG_D_gene | 21 | snoRNA | 12 |
| IG_J_gene | 6 | snRNA | 31 |
| IG_J_pseudogene | 3 | TEC | 28 |
| IG_V_gene | 52 | TR_J_gene | 3 |
| IG_V_pseudogene | 57 | TR_V_gene | 17 |
| lincRNA | 190 | TR_V_pseudogene | 5 |
| miRNA | 43 | transcribed_processed_pseudogene | 9 |
| misc_RNA | 33 | transcribed_unitary_pseudogene | 3 |
| polymorphic_pseudogene | 10 | transcribed_unprocessed_pseudogene | 34 |
| processed_pseudogene | 206 | unitary_pseudogene | 4 |
| processed_transcript | 14 | unprocessed_pseudogene | 155 |

**Table S5.** A detailed biotype list for comparison between mapping methods (Standard - Proposed)

| STANDARD-PROPOSED: Detected only in the proposed method | | | |
| --- | --- | --- | --- |
| **Biotype** | **no.** | **Biotype** | **no.** |
| antisense | 72 | sense_intronic | 9 |
| IG_C_gene | 5 | sense_overlapping | 1 |
| IG_V_gene | 19 | snoRNA | 18 |
| IG_V_pseudogene | 17 | snRNA | 65 |
| lincRNA | 116 | TEC | 11 |
| miRNA | 18 | TR_C_gene | 2 |
| misc_RNA | 134 | TR_J_gene | 5 |
| polymorphic_pseudogene | 6 | TR_V_gene | 35 |
| processed_pseudogene | 603 | TR_V_pseudogene | 5 |
| processed_transcript | 67 | transcribed_processed_pseudogene | 8 |
| protein_coding | 853 | transcribed_unitary_pseudogene | 1 |
| pseudogene | 1 | transcribed_unprocessed_pseudogene | 57 |
| rRNA | 34 | unitary_pseudogene | 2 |
| scaRNA | 1 | unprocessed_pseudogene | 157 |

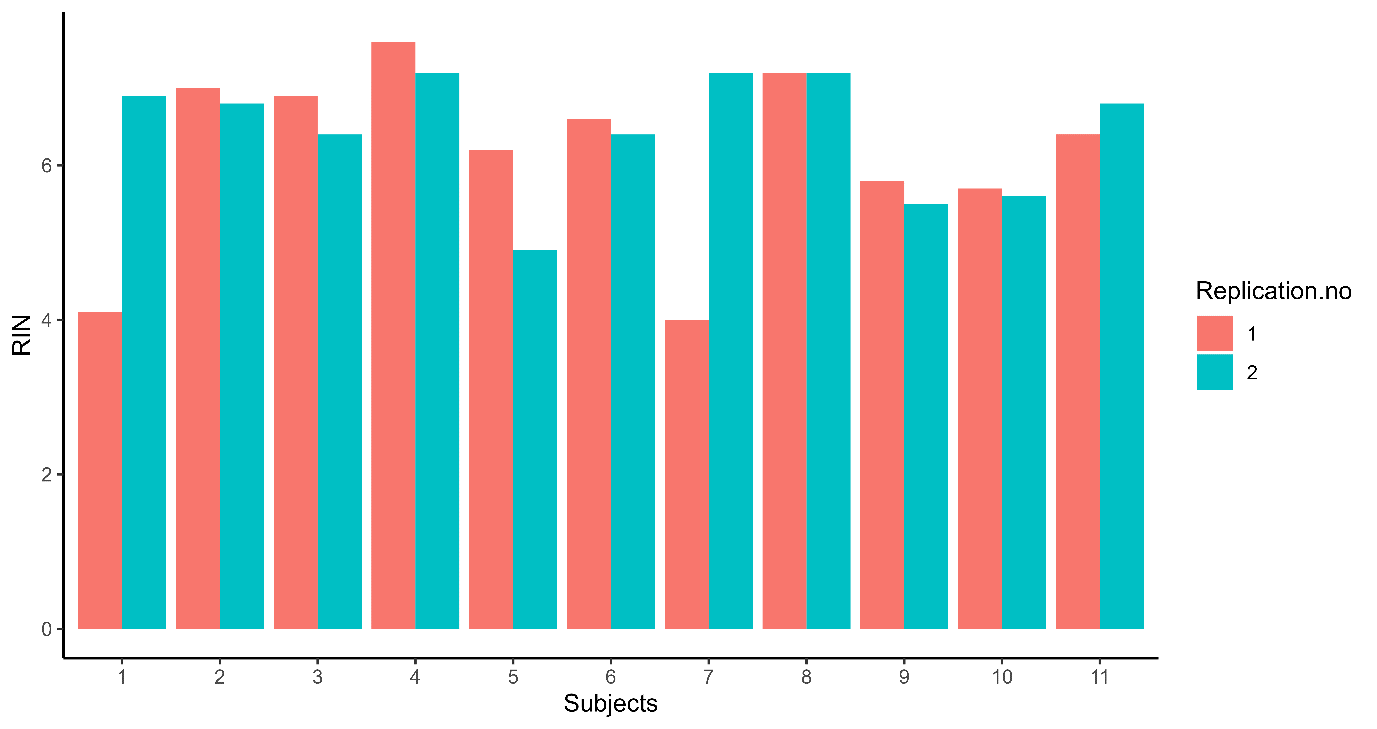

**Supplementary Fig. S1. RIN distribution of replicates**

The x-axis represents subjects, the y-axis represents RIN, and replicates from individual are distinguished by different colors. The RIN ranges from 4 to 7.6 (mean: 6.3, median: 6.5), with RIN differences between replicates from 0 to 3.2.

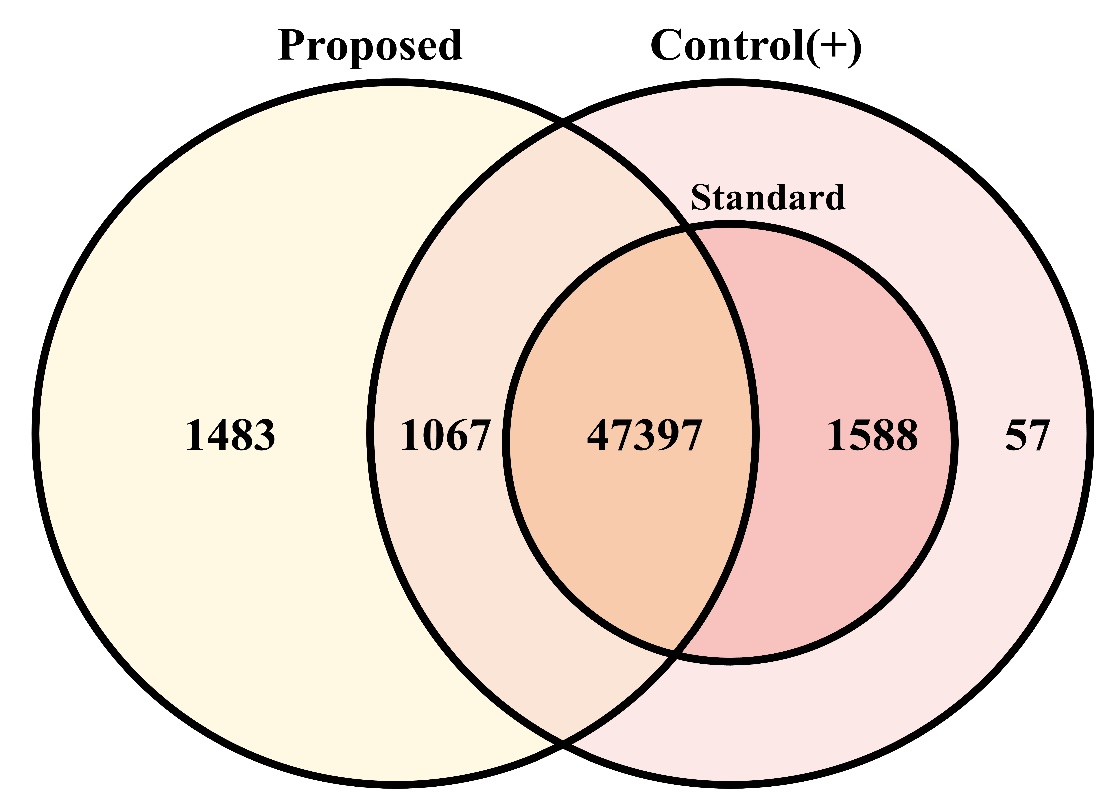

**Supplementary Fig. S2. Number of detected genes among mapping methods**

Venn diagram of genes detected in standard method, proposed method, and positive control.

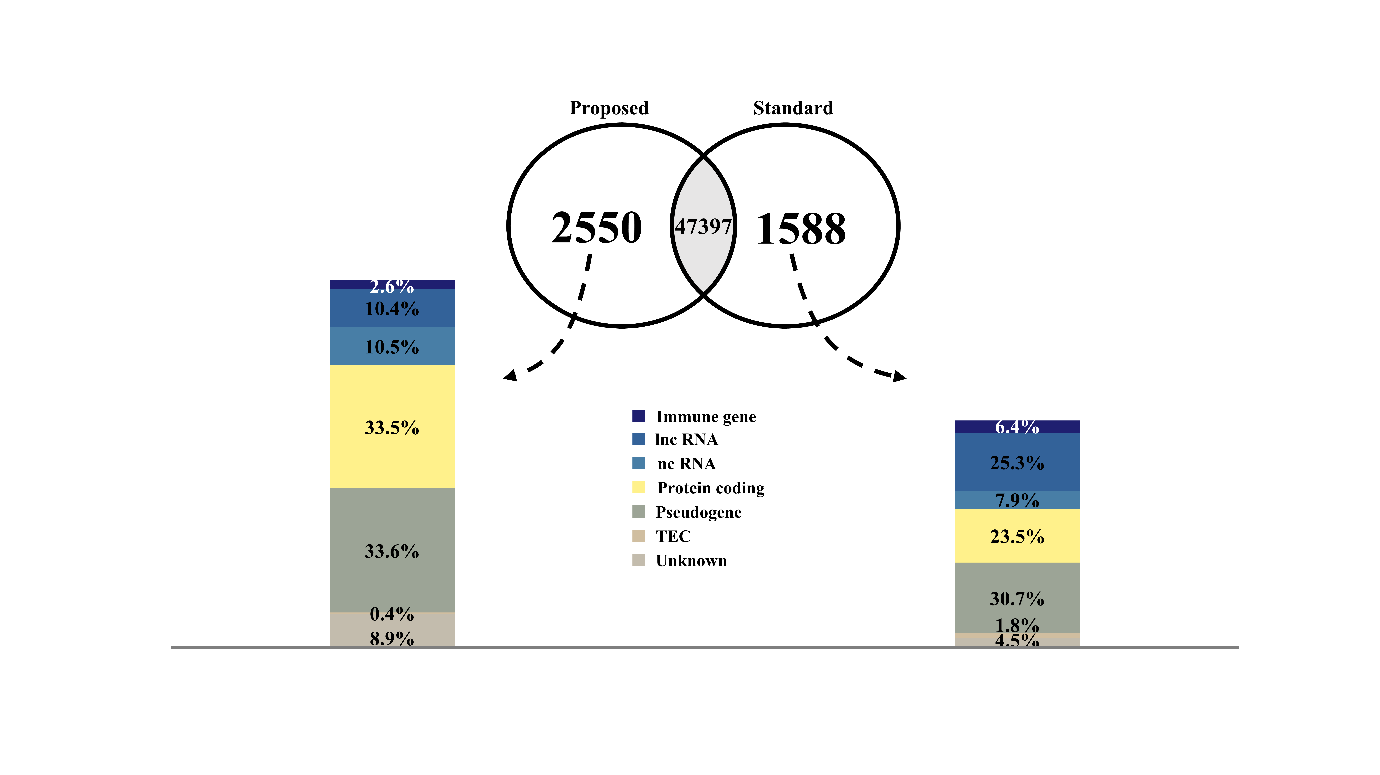

**Supplementary Fig. S3. Biotype comparison between mapping methods**

As a result of comparing g the biotypes of genes detected by the standard method and the proposed method, 2,550 genes were specifically detected in the proposed method and 1,588 genes were specifically detected in the standard method. Biotypes were categorized into 6 types and color-coded, and their numbers were converted into ratios.
